# Generative Machine Learning and Microfluidics uHTS: An Efficient Partnership for Enzyme Engineering

**DOI:** 10.1101/2025.11.02.685536

**Authors:** Pradeep M. Nair, Daniel M. Steinberg, Tiago Resende, Angelica Faith Suarez, Helen Power, Cheng Soon Ong, Vijay Sairam, Shelly Hsiao-Ying Cheng, Lorivie Fragata, Alessandro Serafini, Victor Oh, Say Hwa Tan, Robert E. Speight, Akbar K. Vahidi

## Abstract

Enzyme engineering plays a vital role in tailoring biocatalyst performance to meet the needs of target applications. However, the number of sequence trajectories possible from a single wildtype enzyme sequence is too vast to traverse experimentally. Here we present a novel approach that first expands the experimentally accessible sequence space using ultrahigh throughput screening (uHTS), and then uses indirect and low fidelity assay data to create a “fingerprint” for a target enzyme class. Experimental data from microfluidic uHTS are extracted and used to engineer specificity into an unspecific peroxygenase (UPO) from *Aspergillus brasiliensis* (AbrUPO). We created a library with more than 5 million different variants expressed in *Komagataella phaffii* (*Pichia pastoris*). Microfluidic droplet sorting was then used to generate a dataset of >30,000 unique sequences paired with function data. This dataset was then used to train a task-specific generative model using the Variational Search Distributions (VSD) framework. We compared the variants selected by rank aggregation from the screening data (R series) with novel sequences generated by the refined generative model (G series). While the wildtype enzyme produces nearly equal amounts of both the desired styrene oxide and undesired phenylacetaldehyde products, three out of five of the highest scoring G series variants produced product mixtures more enriched in the desired compound. In comparison, only one of the five highest scoring R series variants showed this improvement. Overall, the variant most enriched in desired product, G929, produced 2.4x more styrene oxide than phenylacetaldehyde, while G3 and G167, produced the highest quantities of desired product at 2.3x enrichment over the undesired product. Further analysis confirmed that our task-specific generative model outperforms existing models pre-trained on large publicly available datasets. This unique combination of uHTS and generative protein modelling provides an intelligent exploration mechanism which not only enables efficient enzyme discovery, but also accelerates optimization and enables predictive insights that are difficult to achieve with either approach alone.

## Introduction

Enzymes are uniquely adaptable catalysts, capable of accelerating reactions by many orders of magnitude with exquisite regio- and stereospecificity.^1–3^ In contrast to heterogeneous catalysts, the reactivity and specificity of an enzyme can be altered through the incorporation of mutations into its genetic sequence. Selecting which mutations to incorporate, however, remains a formidable challenge. The relationship between protein sequence and catalytic function is high-dimensional, non-linear, and dominated by epistatic interactions between residue positions – meaning that beneficial mutations identified individually rarely combine additively.^4,5^ Directed evolution approaches seek to identify beneficial mutation combinations by iterative cycles of mutagenesis to create diverse enzyme libraries, followed by experimental selection to select new backbone enzymes for future cycles. For these workflows to be successful, there are two fundamental challenges that must be overcome. First, the assay used for experimental selection must be predictive of enzymes that will perform best in the final application. Second, enzyme libraries must be constructed that include some candidates with improved performance. These two challenges are inherently coupled; if all possible combinations of mutations were tested for a single enzyme, there is a high probability of finding a beneficial combination. However even a 10-residue peptide contains more than 10 trillion possible combinations of mutations (20^10), far outpacing the throughput of any assay. If library diversity is decreased to be more easily testable, there is a greater likelihood that no beneficial combinations will be present within the library.

Microfluidic droplet sorting provides a platform uniquely well suited to address the first challenge. The ability to compartmentalize biochemical reactions within picolitre droplets, combined with quantitative fluorometric readouts from these droplets, enables fluorescence signals to guide droplet sorting decisions.^6^ Building on this, microfluidic-based fluorescence-activated droplet sorting (FADS) was first demonstrated in 2009 to sort droplets based on enzymatic activity.^7^ This technique was then massively parallelized in ultrahigh throughput screening (uHTS), leading to a roughly a 1,000-fold increase in speed and a 1,000,000-fold reduction in cost compared to traditional automated high throughput screening (HTS).^8^ As a result, microfluidic uHTS dramatically expands the experimental throughput of a directed evolution campaign, enabling millions of enzyme variants to be assayed in a single day. Recent work has made use of such broad datasets to train machine learning models, and then used these models to design improved enzyme libraries for future rounds of directed evolution.^9^ However as a screening platform uHTS is still fundamentally limited to identifying only the best enzymes that are present in a screened library.

Here we propose a conceptual departure from a screening-centered directed evolution platform, taking advantage of the expanded throughput of microfluidic uHTS, without being limited to variants that have been screened. We first use microfluidic uHTS, paired with pooled long-read DNA sequencing, to collect a large dataset of genotype-phenotype relationships for variants of a target enzyme. Rather than using uHTS data exclusively to rank and select variants from within the screened set, we use this coupled dataset as a training corpus for a task-specific generative model. Once trained, the generative model can propose novel sequences that lie entirely outside the screened library – sequences that could not have been identified by direct selection regardless of the depth of the initial screen. In this framing, the uHTS dataset is not merely a hit list: it is a functional fingerprint of the enzyme–substrate system, encoding which sequence patterns correlate with the desired phenotypic outcome across the sampled landscape. The generative model learns this fingerprint and uses it to extrapolate (where appropriate) into unexplored sequence space, proposing variants that have never been physically assayed but that the model predicts will carry the hallmarks of the desired activity. The central claim we demonstrate is that these generative proposals can outperform the best direct selections from the potentially noisy screened data itself.

To connect noisy uHTS data to a generative sequence model in a theoretically grounded way, we employ the Variational Search Distributions (VSD) framework,^10^ a generative Bayesian optimisation approach that learns a conditional sequence distribution over variants with desirable properties. VSD was originally developed for iterative design–test cycles; here we apply it in a single-round setting, using the uHTS corpus as the complete training signal for one cycle of model refinement and generation. The framework explicitly separates two objectives: a discriminative classifier that identifies sequences correlating with the desired phenotype, and a generative model refined using that classifier through a variational inference objective. This separation has important practical consequences. A naïve alternative of fine-tuning the generative model directly on positive examples by maximum likelihood, conflates the two objectives, pushing the model towards reproducing training positives rather than learning the underlying sequence distribution that gives rise to them. VSD’s two-stage procedure instead uses the classifier to provide a smooth, probabilistic signal that guides the generative model towards high-scoring regions of sequence space without overfitting to the finite positive set. A prior model fitted on the entire uHTS dataset simultaneously acts as a regulariser: sequences that score well under the classifier but lie in regions of sequence space unsupported by the observed uHTS data are penalised, guarding against the generation of high-scoring but empirically ungrounded variants. Direct fine-tuning offers no such safeguard; without a prior, the generative model can drift arbitrarily far from the observed sequence distribution if the positive examples pull it there. Crucially, this procedure comes with formal guarantees, ensuring that the generative model is optimised toward the desired functional objective in a principled and convergent manner.^10^

We apply this pipeline to engineer specificity in an unspecific peroxygenase (UPO) from *Aspergillus brasiliensis* (AbrUPO) as a proof-of-concept system. UPOs catalyse H_2_O_2_-dependent oxyfunctionalisation of non-activated carbons, but their promiscuity makes substrate and product specificity an engineering target of practical importance.^11–13^ AbrUPO in particular has been shown to react with at least 16 different aromatic substrates. Catalysis of the reaction between hydrogen peroxide and one of these substrates, styrene, can produce styrene oxide and phenylacetaldehyde. The former is a valuable fine chemical intermediate, so there is industrial value in engineering an enzyme that will catalyse production only of styrene oxide and not of phenylacetaldehyde. We reasoned that directed evolution toward higher substrate specificity would lead to enzymes that catalysed formation of a single product as well. We screened a combinatorial library of ∼5×10^6^ AbrUPO variants by uHTS droplet microfluidics, generating >30,000 unique sequence–function measurements from two parallel assays for substrate depletion. In one assay, we selected AbrUPO variants enriched for styrene conversion, and in a second assay we selected enzymes that showed no reactivity with a different substrate, propylbenzene. These data were used both for direct lead selection (the R series, selected by model-free rank aggregation of uHTS scores) and to train a VSD-LSTM generative model (the G series, sampled from the refined generative distribution). Experimental validation of both series showed that multiple G-series variants outperformed both wildtype AbrUPO and the best direct selections from the screened library, despite the G-series sequences never having appeared in the original screen. The precision-recall performance of the generative model across all 42 experimentally validated outcomes further shows that the VSD-trained model is a meaningful predictor of low-throughput functional outcomes, validating the end-to-end pipeline. Together, these results demonstrate that a single uHTS campaign, combined with task-specific generative modelling, can identify improved enzyme variants that are inaccessible to direct screening selection alone.

## Results

### Combinatorial Library Design

We initially applied two library design concepts to generate diversity in the wildtype that was predicted to improve enzyme performance. First, we used a zero-shot approach to identify residues that are evolutionarily unconstrained and thus more likely to be tolerant to variation. For this purpose, we applied a protein language model (PLM)^14^ to the target sequence. Protein language models provide, for each sequence position, a probability distribution over the 20 amino acids that reflects their compatibility with the learned evolutionary context of the protein family. From this distribution, we extracted the maximum probability per position as a proxy for model confidence and positional constraint. Positions with low maximum probability indicate weak model preference for any specific residue, suggesting reduced evolutionary constraint and, consequently, increased mutational tolerance. Such sites are hypothesised to represent regions of elevated evolvability, where sequence variation can be accommodated without severely compromising structural integrity or function, while still offering opportunities for functional diversification. Based on this rationale, the five positions with the lowest maximum probabilities, D129, L28, D177, F8, and H65, were selected as model-guided mutable sites for targeted exploration (Fig. 1A, red residues). We targeted these sites for combinatorial mutagenesis, to build a mutability sub-library with a theoretical diversity of 2.5 x 10^6^ (SI Methods).

**Figure 1.**
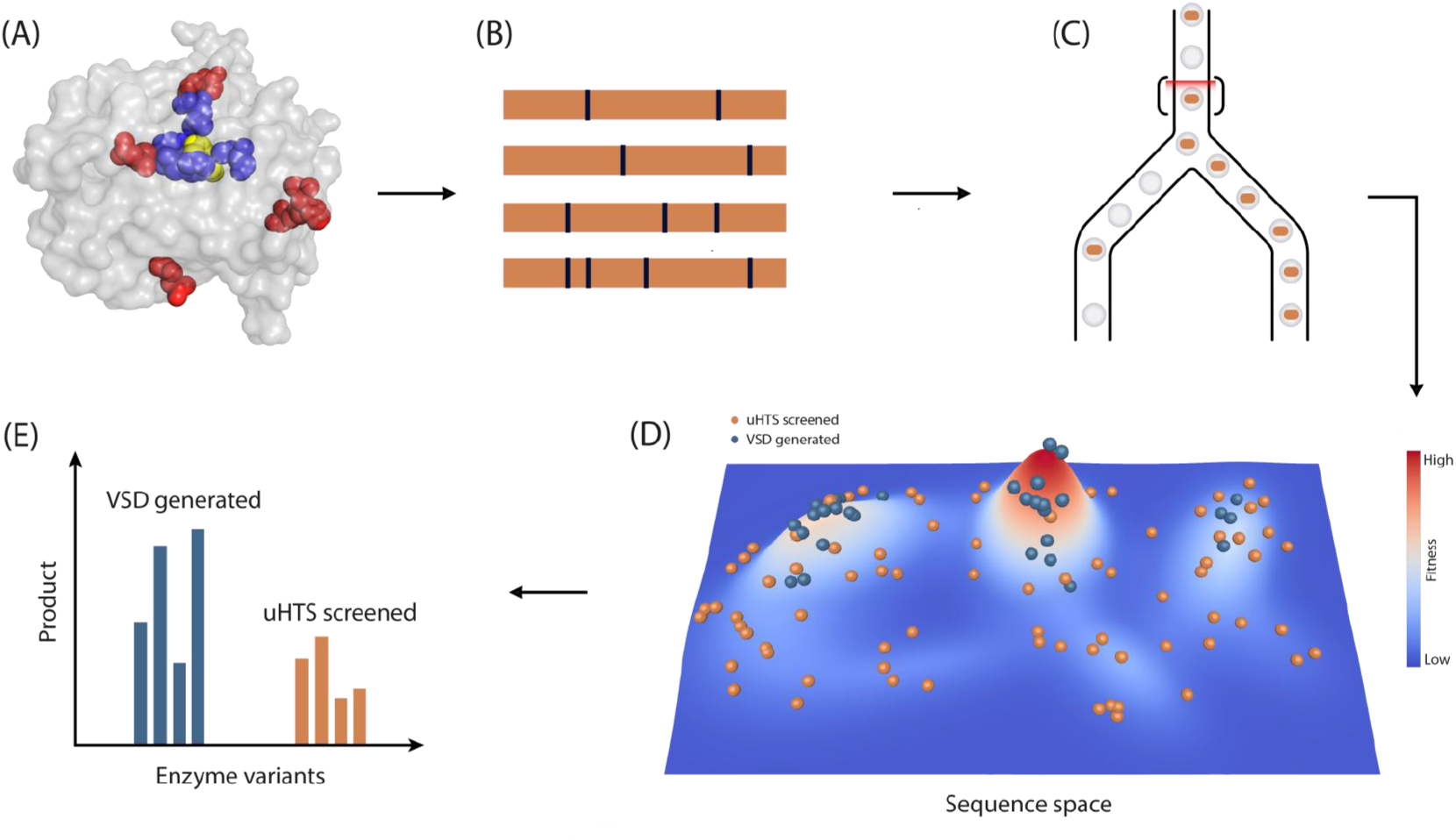
Ultrahigh throughput-guided enzyme modeling. (A) Target residues in the wildtype AbrUPO are selected based on zero-shot predicted mutability (red) and structural active-site residue specificity (blue). The enzyme structure is modeled with an active-site bound styrene molecule (yellow). (B). Gene variants targeting these sites are constructed, then transformed and induced for expression in the host cell. (C) The enzyme library is screened using microfluidic droplet sorting to generate a high volume of data and isolate higher performing variants. (D) Microfluidic droplet sorting data is used to train a task-specific model using the Variational Search Distributions (VSD) framework. Variants predicted to have highest fitness are chosen for experimental comparison to screened variants. (E) Both generated and screened variants are assayed for product formation in parallel.

As our engineering goals were to increase the specific conversion of styrene to styrene oxide, we targeted a second set of residues predicted to be directly involved in substrate recognition. Here, we focused on identifying active site residues directly shaping the styrene binding pocket. A three-dimensional structural model of AbrUPO was generated using protein structure prediction tools,^15^ followed by cavity detection to locate the putative active site.^16^ Docking simulations were then performed with styrene as the target ligand (Figure 1A, yellow structure).^17^ Residues were ranked according to the spatial distance of their side-chain heavy atoms to the docked substrate. The five closest residues, L84, F179, G182, A186, and L231 (Fig. 1A, blue residues), were selected as substrate-proximal positions, under the assumption that mutations at these sites are most likely to alter substrate orientation, binding affinity, or reactivity at the heme center. The substrate binding sub-library was built by saturating these five sites combinatorially, creating a second sub-library also with theoretical diversity of 2.5 x 10^6^.

Together, these ten positions, five chosen by predicted high evolutionary likelihood and five chosen by structural proximity to styrene, were used as the basis for combinatorial library design. The two sub-libraries were pooled together during expression host transformation to build a single library in *K. phaffii* with theoretical diversity of 5.0 x 10^6^, and screening of this library identified 3.1 x 10^4^ unique AbrUPO variant sequences (SI Data1). This dual strategy ensured that the library captured residues predicted to be permissive to sequence variability, as well as positions with direct influence on substrate binding, thereby increasing the likelihood of sampling functional and diverse variants for machine learning–guided enzyme engineering.

### Ultrahigh throughput Library Screening

We designed a microfluidic droplet sorting assay (Figure 2A) in which UPO-expressing cells were grown in droplets and then substrate, either styrene or propylbenzene, and hydrogen peroxide cofactor were picoinjected into the droplets and allowed to react (Fig. 2B). A fluorescent sensor was then picoinjected to convert residual hydrogen peroxide into a fluorescent species. Droplets were then subjected to microfluidic droplet sorting, where droplets were sorted into “Dark” and “Bright” pools based on relative fluorescence intensity (Fig. 3A). More active UPO variants consume the hydrogen peroxide that the fluorescent sensor detects, so these variants are enriched in the dark droplet populations, and inactive variants are enriched in the bright droplet populations. For styrene, out of a sorted library of 144,398 total droplets, those with the lowest 0.6% fluorescence intensity, corresponding to highest UPO activity, were collected in the dark pool (Fig. 3B). For propylbenzene, multiple sorting thresholds were set to create matched pairs of dark and bright pools with varying stringency (Fig. 3C). Propylbenzene sort 1 separated the brightest (most inactive) 95% of 9441 total droplets, sort 2 separated the brightest 99% of 5691 total droplets, and sort 3 separated the brightest 93% of 9724 total droplets.

**Figure 2.**
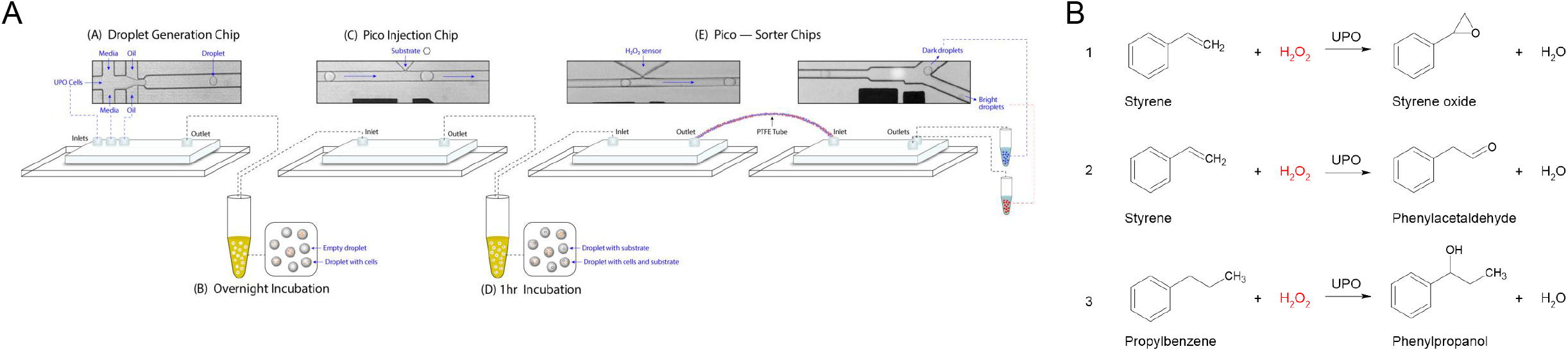
Microfluidic droplet assay. (A) Cells are singly encapsulated in microfluidic droplets in growth media containing methanol for induction of protein expression. Cells are allowed to grow, replicate, and express UPO protein. Substrate and hydrogen peroxide are picoinjected into droplets and UPO is allowed to react. A hydrogen peroxide sensor is then picoinjected into droplets to convert any residual hydrogen peroxide into a fluorescent species. Droplets are sorted by fluorescence intensity. Active UPO will consume hydrogen peroxide, leading to darker droplets. (B) Based on whether styrene or propylbenzene is used as the substrate, reactions 1, 2, or 3 can proceed. In each case, UPO activity leads to consumption of hydrogen peroxide and decreased fluorescence.

**Figure 3.**
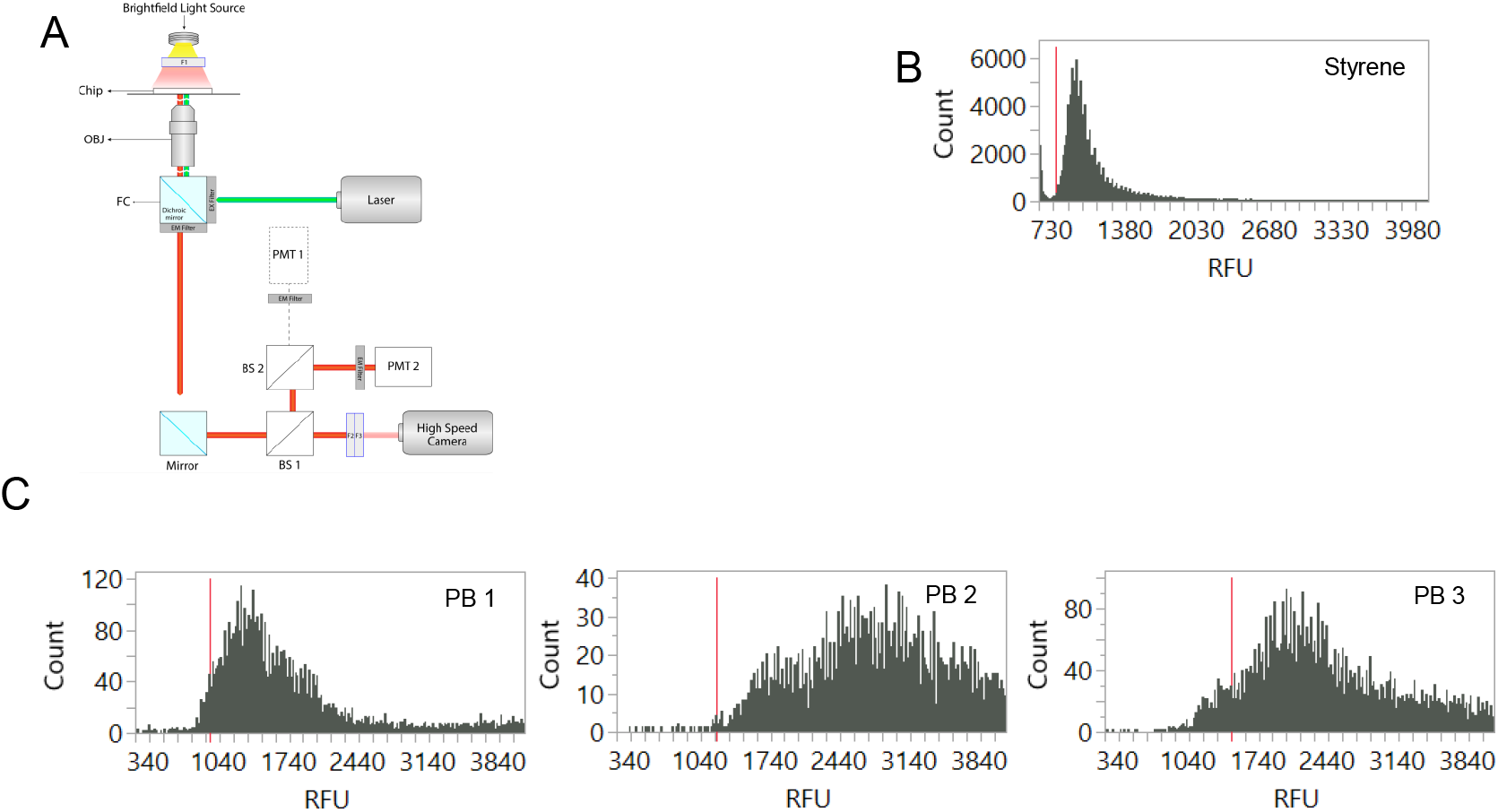
Ultrahigh throughput screening (uHTS). (A) The experimental light path showing excitation with green light, leading to emitted red light that is quantified by PMT 2. (B) Droplet fluorescence histogram from droplet screening for styrene selection. The red line indicates sorting threshold to separate bright and dark droplets. (C) Droplet fluorescence histograms from droplet screening for propylbenzene (PB) selections. The red lines indicate weighted mean sorting thresholds to separate bright and dark droplets.

Pooled, long-read DNA sequencing was then used to determine both the identity and the relative abundance of UPO variants in different droplet populations. We compared the frequency of different variants in different droplet populations, as a function of both substrate (styrene or propylbenzene) and sorting stringency (high, medium, or low for propylbenzene) to score variants in terms of their overall reactivity and specificity for one substrate over the other.

We first used sequencing information directly from uHTS screening to select a subset of variants to build clonally and test for production of styrene oxide and phenylacetaldehyde, as a means of validating our experimental approach (Fig. S1A-D, Table S1). We designed lead identification criteria based on these data (see SI Methods for how leads were selected). Twenty variants that had been screened in uHTS were clonally built into the *K. phaffii* host and induced to express AbrUPO variants. Supernatant was collected from each sample, then used in biotransformation reactions to convert styrene into either styrene oxide or phenylacetaldehyde product. Analytical assays were used to measure the production of the desired compound, styrene oxide, as well as the undesired side product, phenylacetaldehyde. Of these twenty initial variants, ten were selected for re-testing with higher concentrations of styrene substrate to generate a high fidelity validation dataset (SI Data2). To quantify the preference for production of styrene oxide over phenylacetaldehyde we computed a styrene oxide index (SOI) for each variant as follows:

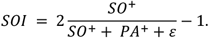

Here SO^+^ = max(0, SO) refers to the amount of styrene oxide product in μM bounded below by 0 μM, and ε is some small amount to guard against dividing by zero (we use 1x10^-5^). This quantity is in the range [-1, 1], where -1 means we have no Styrene oxide product, only Phenylacetaldehyde (PA), whereas 1 means we only have Styrene Oxide (SO). 0 denotes we have an even quantity of both Styrene oxide and Phenylacetaldehyde. From these validation experiments, we identified a single AbrUPO variant, UPO 5, that produced 2-4x more styrene oxide and phenylacetaldehyde (Fig. S2A), though produced more of the undesired phenylacetaldehyde compound, (Fig. S2B). Another variant, UPO 20, displayed a preference for production of styrene oxide compared to phenylacetaldehyde, while still producing > 90% yield of styrene oxide as the wildtype enzyme. Overall, of the 10 tested variants, 5 showed a greater preference for production of styrene oxide than the wildtype enzyme, while 2 showed a greater preference for production of phenylacetaldehyde than the wildtype enzyme. This demonstrated that we were able to enrich for enzymes with the desired product profile using this screening approach.

### Protein Modeling to Generate Novel Diversity

Having validated that the uHTS approach did identify variants with greater activity or specificity for the desired compound, we next asked whether the high volume of screening data focused on only the initial biocatalytic reactions of interest could be used to train a machine learning model to predict previously untested variants with desired reactivity profiles. For this task, we built multiple generative models trained with the uHTS dataset (summarized in Table S2, and visualized in Fig. 4). We then tested these models for their correlation to the high fidelity, low volume validation dataset with direct uHTS leads (SI Data2). These test results were used to select the highest performing generative model for subsequent variant generation in the 10 pre-selected sites. We found that an LSTM generative model yielded the best performance on the initial low volume, high fidelity validation dataset (Table S3, Fig. S3). This LSTM model was trained using a classifier scorer, as opposed to directly learning from the uHTS data. The classifier was itself trained to predict highly active and substrate specific sequences from the uHTS dataset. We detail how we created classifier targets in the methodology. This classifier scorer allows the LSTM to extrapolate beyond the uHTS dataset using the variational search distributions (VSD) algorithm.^10^ For a baseline to compare the generative model to, we used the uHTS data directly as a “model free” variant selection method. We summarize the two compared selection methods as:

**Figure 4.**
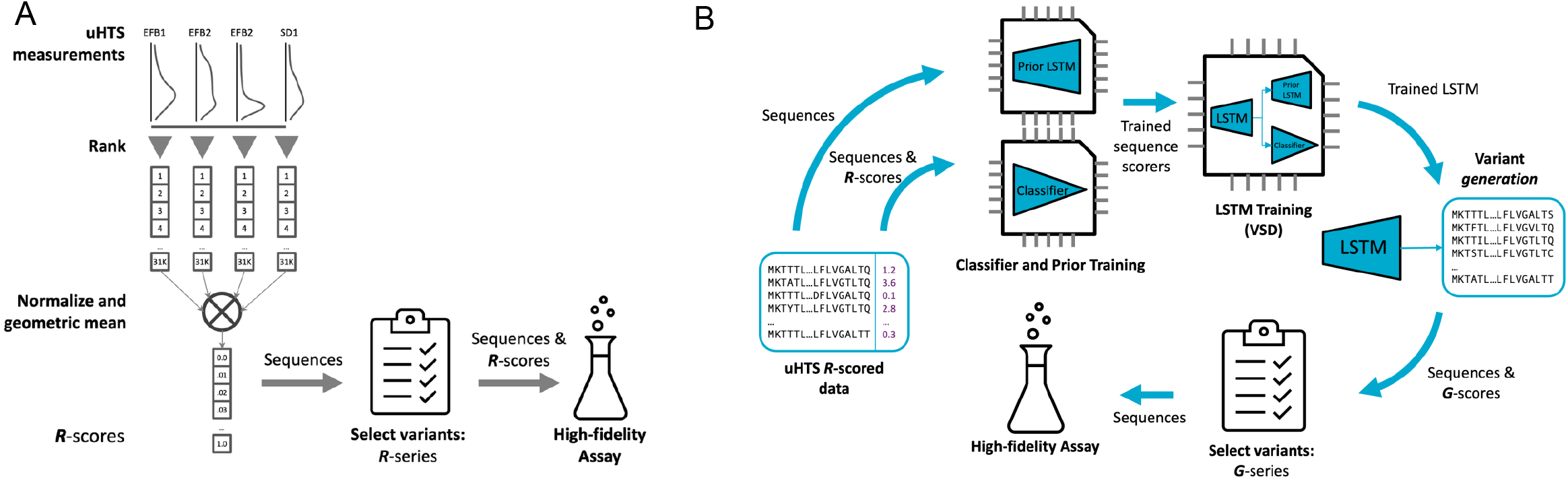
Model training data from uHTS. (A) Visualisation of the rank aggregation process to compute R-scores. (B) Visualisation of the VSD LSTM training process and computation of G-scores.

***R*** – direct selection from 30,801 variants appearing in the microfluidics sequencing data that had more than one styrene “dark” read, using an aggregated ranking score^18^ (R-score) of preference for styrene substrate specificity. See Figure 4A, and SI Methods for details.

***G*** – selection from 4,374 *entirely novel* variants generated by a task specific LSTM, trained using the VSD algorithm and the R-scores. The generated variants were scored using the log-likelihood of the generative model (G-scores), and generated variants that also appeared in the uHTS data were removed. See Figure 4B and SI Methods for details.

For experimental testing, we chose the top 5 scored variants from each method, as well as 5 logarithmically spaced scored variants from each method, i.e. 10 selections per method. The top 5 variants reflect those that would typically be chosen for further study in an engineering campaign. The logarithmically spaced variants are meant to provide a very sparse picture of the success of either approach in identifying improved variants. We refer to the variant scoring methodologies as R-scores and G-scores respectively, and the final selected variants as the R-series and G-series respectively – see Tables S4 and S5 for the selected variants. To differentiate between the initial selection pools, we refer to the 30,801 sequences from uHTS screening with styrene substrate as R, and the 4,374 LSTM generated variants as G. This approach yielded a total of 20 variants to be tested, in addition to wildtype AbrUPO. These variants were clonally built and expressed in the *K. phaffii* host, then assayed for production of styrene oxide and phenylacetaldehyde, as described previously (Fig. 5A). Several of the supernatant samples showed no conversion to either product. Western blotting confirmed protein expression upon induction for all transformants (Fig. S4A-C).^19^ We cannot assess whether unreactive samples were unreactive because of inactive enzyme or poor enzyme expression or folding. Among those samples that did show reactivity, we observed that two of the G-series, G3 and G167, produced 2-2.3x more of the target styrene oxide product, and produced 2.3x more styrene oxide per mole of phenylacetaldehyde, compared to the wildtype AbrUPO (Fig. 5A). A third member of the G-series, G929, produced roughly half as much styrene oxide as wildtype AbrUPO, but produced 2.4x more styrene oxide per mole of phenylacetaldehyde compared to wildtype AbrUPO. In contrast, only R10K from the R-series was improved along either metric relative to AbrUPO, producing significantly more styrene oxide per mole of phenylacetaldehyde, ∼2x higher than wildtype AbrUPO.

**Figure 5.**
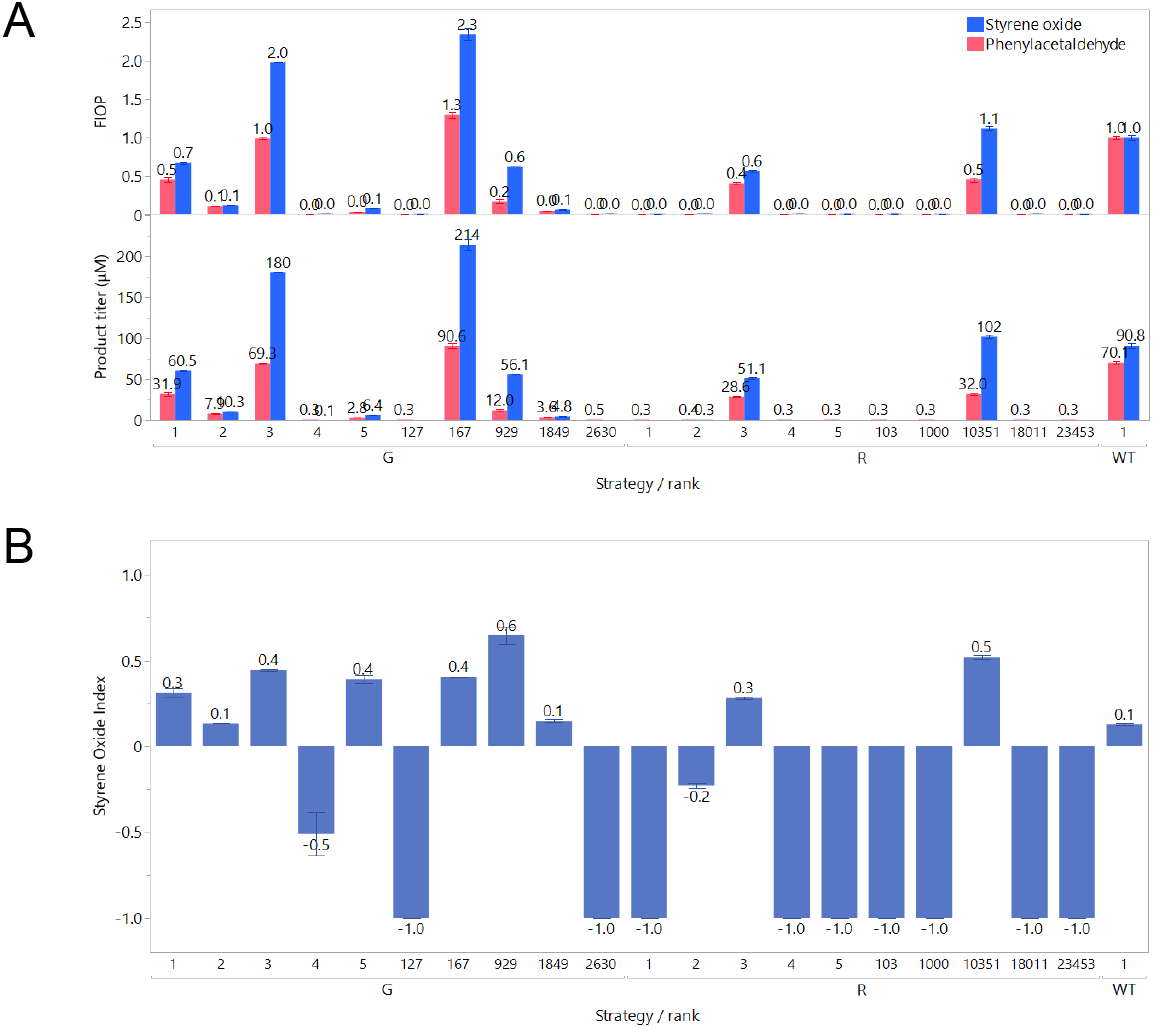
G- and R-series performance. (A) For G- and R-series UPO variants: Productivity of styrene oxide and phenylacetaldehyde, measured by gas chromatography. Fold increase over parent (FIOP) was calculated by normalizing product titers of each UPO variant by that of WT UPO. (B) Styrene oxide Index (SOI) was calculated for each variant. An SOI of 1 corresponds to formation of only styrene oxide and an index of -1 corresponds to only phenylacetaldehyde. Error bars denote standard deviation across duplicate biotransformation reactions.

To quantitatively assess the ensemble differences between the R-series and the G-series, we compared the top 5 selected variants from both strategies, as well as a matched ranking from both strategies, G3 and R3, in Table 1. We found that in most cases, the generated variants, G, were significantly (α = 0.01) better performing than the aggregated rank variants, R, using a Brunner Munzel^20^ test with an alternative hypothesis of *p*(*y*_*G*_ > *y*_*R*_ > 1/2. The top 5 selected variants demonstrate the difference in the predicted best performing variants under either strategy. This subset also emulates how one would typically use the R- and G-scores to select variants for testing, e.g. one would select the top-k scored variants.

**Table 1.** Average experimental results for the **top 5** selected variants from the G and R methods, as well as a matched ranking from both methods, and Brunner Munzel test p-values under the alternative hypothesis that the distribution of G is stochastically greater than that of R.

| Strategy | Styrene oxide (μM) | Phenylacetaldehyde (μM) | Samples (N) |
| --- | --- | --- | --- |
| **G mean** | 51.48 | 22.44 | 10 |
| **R mean** | 10.07 | 6.00 | 10 |
| **WT** | 91 | 70.1 | 2 |
| **Best (G3)** | 180 | 69.3 | 2 |
| **Best (R3)** | 51 | 28.6 | 2 |
| **Brunner**<br>**Munzel**<br>**p-value** | 0.0011 | 0.0091 | 20 |

In the VSD framework, the LSTM was trained in two stages; firstly on the microfluidics sequences only (no experimental outcomes), and secondly it was refined using a classifier trained to discriminate the top 10% of R-scored sequences to score sequences generated by the LSTM. We refer to these as the “prior LSTM” and “VSD LSTM”, respectively. The function of the prior LSTM is to constrain the VSD LSTM generated sequences to not be too different from those seen in the microfluidics sequencing data, and the function of the classifier is to push the VSD LSTM to generate predicted high-performing sequences, possibly *outside* the set of seen sequences.

We also sought to more thoroughly understand the VSD LSTM’s selection performance by testing the model’s ability to select “useful” variants from the full 42 experimental outcomes on two binarized target criteria, with the proportion of positive examples:

1. Styrene Oxide > WT (92.87), Proportion positive labels: 0.143
2. Styrene oxide index > WT (0.13), Proportion positive labels: 0.429

The first target captures the engineering goal of increased styrene oxide production. The second target is a stringent criteria that identify variants more specific than the wildtype enzyme to produce styrene oxide compared to phenylacetaldehyde.We used the ([0, 1]-normalized) log-likelihoods of the generative model (G-scores) as the criterion to select variants by thresholding, then measured how many positive and negative variants according to the preceding 2 criteria were selected for different threshold levels. We summarize results as precision-recall curves in Figure S5, where we use bootstrap sampling (with 1000 bootstrap samples) for creating intervals. Precision can also be understood as the true discovery rate (true positives / true and false positive) and recall the sensitivity (true positives / true positives and false negatives). When choosing a threshold on the log-likelihood for selection, precision-recall curves can help us understand the trade-offs we face, as choice of threshold will directly affect precision and recall outcomes. For example, if we chose a low threshold so that all variants are selected, then we would capture all the positive variants, maximizing recall, but we would also select all the negative variants (as false positives), lowering precision. Conversely, if we set the threshold so that only the highest log-likelihood variant is selected, then we will achieve perfect precision if this variant is positive, but low recall because we miss all the other positive variants in the set (false negatives). We can also summarize these curves by measuring the average precision (AP) for all recall values in the curve. An AP of 1 means perfect precision and recall, and an AP that is the same as the chance level of precision means the predictor is no better than random.

We summarize the APs below (with 95% confidence interval in brackets),

1. Styrene Oxide > WT: AP = 0.244 (0.183, 0.458)

2. Styrene Oxide Index > WT: AP = 0.602 (0.483, 0.813)’

All APs are better than chance (including the lower confidence limit). We found that the VSD LSTM performed significantly better than chance (randomly selecting variants according to the base positive label rate) for all recall levels, in all cases. The VSD LSTM maintained high precision for all recall levels for Styrene Oxide Index > WT criteria, but Styrene Oxide > WT has a lower precision, most likely because the wildtype sequence already exhibits high production for styrene oxide, resulting in fewer positive examples. Note that since some of these variants were chosen using the VSD LSTM (particularly the top-5 G series), there may be some slight selection bias present in these results, however we believe that these plots are still useful in indicating how useful the VSD LSTM would be if it was the only method used to select variants for experimentation, and a threshold selection criterion was used as opposed to our top 5 methodology.

Protein language models (PLMs) have recently demonstrated utility for predicting protein properties of interest. Model families such as ESM1/2,^21^ ESMC^22^ and ProtTrans^23^ have been trained on datasets of up to 6.8 billion sequences. They learn the evolutionary distribution of protein sequences, capturing the conservation and co-variation patterns that reflect selective pressures spanning billions of years and diverse protein families. Without additional training, PLMs reflect evolutionary plausibility and were therefore used to in the early stages of this study to identify suitable sites for combinatorial mutagenesis. However, applying PLMs to task specific predictions such as enzyme-substrate specificity requires additional training on large task specific datasets.

To place our *task-specific generative* VSD LSTM in context, we benchmarked it against relevant PLM-based models for the enzyme-engineering task explored in this study. Models were chosen based on a combination of their trained tasks and reported performance metrics. We evaluated each pretrained supervised model alongside the VSD LSTM and the prior LSTM using Spearman correlation against all experimental outcomes. We used the predicted values from the pretrained models and the likelihood of the sequences under the generative models for the predicted scores, with results summarized in Table 2 and SI Methods. See SI Data3 for the complete tested set. Note that for Boltz unconstrained affinity scores, a lower value means a higher predicted affinity for the substrate and therefore negative Spearman’s correlations indicate a stronger relationship. We only show the highest correlated pretrained models. While both phenylacetaldehyde and styrene oxide production did correlate with the listed model scores, in all cases the VSD LSTM model had the highest correlation. The prior LSTM also performs comparatively well even though it has only seen sequences and microfluidics measurements. This relatively high performance is because most sequences detected by the uHTS process were taken from the styrene-active pool of variants. We must caution however, that these results may suffer from some selection bias, as VSD LSTM was used to select almost half of the samples. Furthermore, these results should be viewed in light of the fact that the pretrained supervised models cannot be used to generate novel sequences directly, unlike the VSD and prior LSTM models. They can only be used for in-silico “screening” on target sequences that have been explicitly defined *a priori*. Such a set of hypothesised sequences must therefore already exist before pre-trained models can be applied.

**Table 2.** Spearman correlations and p-values (in parentheses) between the generative and pretrained models on the 42 experimental samples (duplicate samples of 10 R-series, 10 G-series, and AbrUPO wildtype).

| Assay | VSD LSTM | Prior LSTM | NetSolP solubility | Boltz unconstrained binding probability | Boltz unconstrained affinity (more negative is stronger relationship, see text) |
| --- | --- | --- | --- | --- | --- |
| Phenylacetaldehyde ($\mu\text{M}$) | **0.447 (0.003)** | 0.360 (0.019) | 0.392 (0.010) | 0.163 (0.303) | -0.154 (0.331) |
| Styrene oxide ($\mu\text{M}$) | **0.443 (0.003)** | 0.363 (0.018) | 0.416 (0.006) | 0.146 (0.357) | -0.308 (0.047) |

## Conclusion

Engineering enzymes to achieve higher production of target chemicals remains a central challenge for biocatalysis. In this work, we demonstrated that uHTS data can provide high volume, low fidelity measurements sufficient for effectively training task-specific generative models. The utility of uHTS in providing a training dataset also provides a template for how best to utilize the high volume of uHTS assay data without requiring a fully tailored assay. Here, we used a uHTS assay that detected depletion of a conserved substrate for multiple reactions. The ability to use a modular assay greatly broadens the applicability of this paired uHTS-generative machine learning approach for enzyme engineering. Using the Variational Search Distributions (VSD) framework to refine a long short-term memory (LSTM) model on these uHTS data, we were able to generate novel *Aspergillus brasiliensis* UPO variants with improved performance compared not only to the wildtype enzyme, but also to the best variants directly identified from uHTS selection. Several generated sequences achieved substantially higher styrene oxide yields and specificity scores, validating that low-fidelity measurements can be effectively leveraged to guide design toward higher-fidelity experimental outcomes.

Our findings underscore three key insights. First, low fidelity screening data, whilst noisy and indirect, are nonetheless valuable for informing generative models that can extrapolate beyond the training set to propose superior variants. Second, a relatively simple generative architecture, when refined via VSD conditioning, consistently outperformed direct selection from the same screening data— illustrating how generative modeling can transcend the limits of the assay used for training. Third, while some pretrained protein language models achieved moderate correlations with assay outcomes, their predictive value was inconsistent or hard to arrive at *ex ante*, emphasizing the advantage of models explicitly trained on task-specific, experimentally grounded data.

Overall, this study illustrates a generalizable strategy: breadth of coverage through ultrahigh throughput, low fidelity screening, coupled with generative modeling to translate that data into high fidelity design gains. Armed with a model trained on an enzyme class of interest, it is now possible to proceed to intelligently designed specific variants without the onerous task of multiple rounds of deep mutational scanning. By exploiting the complementary strengths of scale and extrapolation, this approach accelerates the discovery of enzyme variants with improved production of specific compounds, and offers a blueprint for applying machine learning to protein engineering problems where conventional data collection is limited.

## Acknowledgements

D.M.S., H.P., C.S.O. and R.S. were funded by the CSIRO Advanced Engineering Biology Future Science Platform and Science Digital Program, and were supported by resources and expertise provided by CSIRO IMT Scientific Computing. For helpful discussions and support of this work, we thank Michael Burns, Audrey Robic, Jayer Tan, and Muhummed Khabir. We thank Charmaine Lee for graphical depictions of the project workflow.

## Contributions

Conceptualization, P.M.N., D.M.S.; Library design, T.R., D.S., and P.M.N.; Cloning S.H.-Y.C., L.F., A.F.S., A.S., and P.M.N.; Microfluidic screening, S.H.T., V.S., V.O.; Analytical chemistry, A.F.S.; Sequencing, S.H.-Y.C., A.S., A.F.S., T.R.; Project supervision, A.K.V., and R.S.; Protein modeling, D.S., H.P., T.R., and C.S.O.

## Declaration of interests

Authors P. M. N., A. F. S., S. H. T., V. S., V. O., S. H.-Y. C., L. F., T. R., and A. K. V. are inventors on patent application 2504125.2, that describes the ultrahigh throughput screening platform used here, and improved enzyme variants identified using that platform.

## Notes

### Summary of Updates

Experimental data using HPLC to measure phenylpropanol and styrene oxide has been removed to focus on highest fidelity data, GC measurements of styrene oxide and phenylacetaldehyde production; A new metric for styrene oxide production over phenylacetaldyide, SOI, has been calculated and is compared in figures 5, S2, and S5; Panels in figures 1-3 have been edited for clarity; Text has been edited for clarity and focus.

