## Supplemental Methods and Tables S1-S5 for "Generative Machine Learning and Microfluidics uHTS: An Efficient Partnership for Enzyme Engineering"

***K. phaffii* cloning**

An expression host was generated by integrating *ADE2* gene into *PichiaPink* Strain 4 (Δade2, Δprb1, Δpep4; Invitrogen) at *AOX1* locus. The *ADE2* gene was driven by the pPADE2_LC promoter, PCR-amplified from the pPINK-LC plasmid. The resulting strain, STR1354, was used for expression of AbrUPO libraries and variants. An expression cassette was synthesized (GenScript, Piscataway, NJ, USA) with the following parts:

- AbrUPO gene (NCBI OJJ73116.1), codon-optimized for *K. phaffii*, or AbrUPO enzyme variants identified through screening or modeling approaches, fused with a C-terminal His-tag
- TDH3 terminator (TDH3tt), amplified from the *PichiaPink* Strain 4 genome
- Zeocin resistance marker, amplified from pPICZA, inserted downstream of TDH3tt
- Complete AbrUPO–TDH3tt–Zeo cassette behind an AOX1 promoter and flanked by P1NS4 homology arms to enable genomic integration and methanol-inducible expression.
- The plasmid backbone containing an ampicillin resistance marker and f1 origin of replication, was amplified from pBluescript II SK(+)

The plasmid was propagated in *E. coli* NEB-5α (New England Biolabs) and used as a template for AbrUPO variant generation. For transformation into STR1354, the expression cassette was linearized and amplified by PCR with the following primers:

- SHC231: GCCTTGGACATCGACCC
- SHC244: TGTTGGTATACATTTCCTTATCCATTAG

The PCR product was purified using the Zymo Clean & Concentrator kit, and 1 µg of DNA was used for transformation. Electroporation-competent *K. phaffii* cells were prepared according to the *PichiaPink Expression System Manual*. Following electroporation, cells were recovered in YPDS medium for 4 hours and either plated or grown in liquid culture containing YPD and 100 µg/mL zeocin.

**Library generation**

To build diverse libraries of *K. phaffii* strains, the expression cassette plasmid was mutagenized with site-directed mutagenesis (QuikChange, Agilent, Santa Clara, CA, USA). Two degenerate codon sets were used for transformation, NDC and VHG, which collectively encode 19/20 naturally occurring amino acids and no stop codons. Each sub-library (mutability and substrate binding) was prepared separately by mutagenizing the AbrUPO expression plasmid using five iterations of QuikChange. Mutagenized expression cassettes were PCR-amplified and then the two sub-libraries were combined and used for a single *K. phaffii* transformation as described above. Following electroporation and recovery in YPDS medium, libraries were separated into fractions for plating on solid agar and growth in 600 µL YPD media supplemented with 100 µg/mL zeocin in 96 well deep well plates. Agar plates were incubated for 3 days at 30 $^{o}$C and liquid cultures were incubated for 3 days, at 30 $^{o}$C, with 80% humidity and shaking at 980 RPM in a Multitron plate incubator (Infors HT). After growth, library liquid cultures were harvested directly for screening or combined with 50% glycerol solutions to prepare glycerol stocks (25% glycerol final concentration) for long term storage at -80 $^{o}$C.

### Microfluidics

The microfluidic devices used in the experiments consist of a droplet generation module[28], a pico-injection module[29] and a droplet sorting module[30]. The devices are fabricated in Poly(DiMethylSiloxane) PDMS (Dow Corning) using standard photo-lithography and soft lithography[31]. All the devices are treated with Aquapel (PGW Auto Glass) to render the surfaces hydrophobic. The electrodes in both the pico-injection module and droplet sorting module were produced using the microsolidics[32] technique. In the pico-injection module, the electrodes are placed directly opposite to the pico-injector. In the sorting module, the electrodes are placed directly below the sorted droplet channel.

The droplet generation module used a double cross junction[28] to create monodispersed water-in-oil (W/O) droplets. The diameters of the droplets are about 35 μm with less than 5% standard deviation in size. In this configuration, two aqueous phases (Aqueous Solution A containing cells, growth media, zeocin, and methanol and Aqueous Solution B containing only the growth the media, zeocin, and methanol) were encapsulated with a single continuous oil phase (Novec 7500, 3M) to form W/O droplets. These droplets were stabilized using surfactant that prevents droplets from merging and ensures that the droplets are stable over long periods of time (more than 3 days) with minimal material transfer between the droplets. The W/O droplets contain both the *K. phaffii* cells and induction media within a controlled microenvironment so that the cells are contained within the droplets. In our experiments, single cell encapsulation follows a Poisson distribution[33] which is about 10%, verified using image analysis. The fluorinated oil is well-known to support cell growth due to its high oxygen solubility and inert properties.[34] After droplet generation, we incubated the droplets at 20 $^{o}$C for 2 days to allow the cells to grow within the droplets.

After incubation, the W/O droplets were reinjected into a pico-injection module to allow the substrate solution (Solution C, containing substrate and $H_2O_2$) to be pico-injected into the droplets. The pico-injection rate was about 1200 droplets per second (dps) and the volume injected was about 1.5 pL. We collected about 3 million droplets and further incubated the droplets for an hour at 25 $^{o}$C to allow UPO variants to react with the pico-injected substrate.

After the one-hour incubation, we reinjected the droplets into a second pico-injection module connected in serial to a sorting module using polytetrafluoroethylene (PTFE) tubing. This connection allowed us to maintain the required incubation time. Here, about 2.5 pL of Solution D, containing a fluorescent sensor for $H_2O_2$, was pico-injected into each droplet. The droplets were then sorted into different collection tubes based on their fluorescent intensities.

### Clonal expression and harvest

Beginning with clonal patches on solid agar, colonies were picked into individual wells in 96 well deep well plates in BMG minimal media supplemented with 100 µg/mL zeocin. Strains were grown for 2 days at 30 $^{o}$C, with 80% humidity and shaking at 980 RPM in a Multitron plate incubator. Strains were then induced by the addition of methanol and grown for 24 hours more at 20 $^{o}$C, with 80% humidity and shaking at 980 RPM. Supernatants were then collected for all wells, then pooled by strain.

Supernatants were concentrated using Corning Spin-X® UF concentrators (membrane cutoff of 10k Da), with 6 or 20 mL volume capacity depending on the volume of culture. All supernatants were buffer-exchanged with 100 mM potassium phosphate buffer pH 7.0. Protein concentrations were measured using the Qubit Protein assay kit (Thermofisher, 33212).

### Biotransformation of substrates

Reactions were conducted in a 400 μL volume in 2.0 mL reaction tubes at 25 ℃ and 600 rpm in duplicate. The reaction mixture contained the UPO strain supernatant, 10 mM of substrate (dissolved in DMSO), 4 mM of $H_2O_2$, 2 mM of $MgCl_2$. All reactions were run in 100 mM potassium phosphate buffer, pH 7.0. After overnight incubation, reactions were processed for GC analysis.

**GC analysis**

Styrene and its products (phenylacetaldehyde and styrene oxide) were detected using GC analyses. Ethyl acetate (200 μL) was used to extract styrene substrate and products from the biotransformation mixture and 1 µL of the sample was injected in split mode (5:1). The following conditions were used: GC column HP-5 30 m x 0.320 mm x 0.25 µm, carrier gas: He, 1 mL/min; injector temperature: 240 ℃; detector temperature: 300 ℃; oven temperature: 60 ℃ (initial time 2 min), rate, 20 ℃/min, final temperature, 300 ℃ with total run time of 16 min. Peaks were assigned by comparison to chemical standards prepared from commercial styrene, phenylacetaldehyde, and styrene oxide. Using this method, styrene eluted at 4.2 min, phenylacetaldehyde at 6.3 min, and styrene oxide at 6.5 min.

**Validation of uHTS**

We selected UPO variants to build clonally and test for production of styrene oxide and phenylacetaldehye. The selection criteria for which clones to build included 2 factors:

1. Abundance in the active (dark) fractions of the styrene sorted population

2. Enrichment factors or ratios of variants in the inactive (bright) propylbenzene fractions of propylbenzene sorted fractions

Four different selection criteria concepts were applied to the variants obtained in the uHTS. A total of twenty variants (5 variants per concept) were selected using different combinations of these factors. The concepts focused on ranking variants based on higher enrichment factor of different stringency propylbenzene pools (higher to lower) (concept 1), then ranking variants based on overall enrichment factors (concept 2), then ranking variants based on variant sequencing read counts in the styrene pool (concept 3) and finally ranking variants based on higher read count ratios of different stringency propylbenzene pools (higher to lower) (concept 4).

UPO variants were substituted in place of wildtype UPO in the expression construct described above, then plasmids with each variant expression vector were synthesized. Expression constructs were linearized and amplified by PCR, then cloned into *K. phaffii* as described above.

Clones of each variant were assayed for activity, using gas chromatography to detect styrene oxide and phenylacetaldehyde. Their sequences are shown in Table S1. Those that showed activity were re-screened using higher substrate concentrations to gain greater confidence in their activities. The higher confidence re-test values are reported.

**Model free rank aggregation: R-scores**

The “R” series refers to the model free rank aggregation method, which is purely based on ultrahigh throughput microfluidics measurements. We have four “low fidelity” targets from the microfluidics measurements that proximally measure substrate specificity when combined:

1. EFB1: Enrichment factor threshold 1 – propylbenzene substrate
2. EFB2: Enrichment factor threshold 2 – propylbenzene substrate
3. EFB3: Enrichment factor threshold 3 – propylbenzene substrate
4. SD1: Styrene dark read count – styrene substrate

For styrene substrate, a single pool of dark droplets was collected. Data from this pool was in the form of sequencing read counts in the dark pool. Since dark droplets have less residual hydrogen peroxide, these droplets should have more active UPO variants. So styrene dark read count is expected to be correlated with peroxygenase activity.

For propylbenzene substrate, three screening thresholds were used. At each threshold, both dark and bright pools of droplets were collected, and an enrichment factor was calculated for each variant in one of the pools collected at each threshold. Enrichment factor is computed as:

$$EF = \frac{Dark\ read\ fraction}{Dark\ read\ fraction\ +\ Bright\ read\ fraction}$$

And so is anti-correlated with peroxygenase activity. For each of these targets, we obtained the number of variants in Table S2, where we only count variants with more than one sequencing read. We also plot histograms of these quantities in Figure S1.

We used a rank aggregation method[25] to aggregate these separate targets into a single "low fidelity", high volume, proxy for substrate specificity target, $R \in (0, 1)$.The aggregation algorithm is as follows,

1. For each target vector, $t \in \{EFB1, EFB2, EFB3, SD1\}$,
    a. $r_t \leftarrow rank(y_t)$, where $r_t = 1$ is best and corresponds to $\max(y_t)$, missing values are ranked last, otherwise ties are given an average rank.
    b. $\tilde{r}_t \leftarrow \frac{r_t}{\max(r_t+1)}$ to obtain a normalized score, $\tilde{r} \in (0, 1)$

Where $y$ denotes microfluidics measurements.

2. Take the geometric mean of the ranks as the aggregated score,

$$R = \exp\left(\frac{1}{4}\sum_t log(r_t)\right)$$

This rank aggregation score, R, is maximally correlated with the original rank orderings of the low fidelity targets, and therefore should score variants with high EFs (low propylbenzene activity) and high SD1 (high styrene activity) well (low R). Conversely, low (or missing) EFs, and low SD1 are ranked poorly (high R). See Figure 4A for a visualization of the R-score calculation. The top 5, and then 5 logarithmically spaced variants were chosen according to their R score – see Table  for the final list.

### Variant generation and scoring using generative models: G-scores

The "G" series refers to the generative model method, which is informed from the aggregated ranks, "R". We use Variational Search Distributions (VSD)[16] for training a generative model on the microfluidics data. VSD is a solution to the active generation problem, where we want to actively learn a conditional generative model from experimental data, such that it can learn to generate designs (e.g. proteins) that are optimal according to the experimental outcomes. Typically, the problem is specified for multiple rounds of experimentation, but here we run just one round. VSD is comprised of three components. The first is a prior generative model:

Prior: $p_\psi(\boldsymbol{x})$.

Where $\psi$ are the parameters of the generative model, and $\boldsymbol{x}$ are (integer encoded) sequences. The function of the prior is to define the initial search space. The next component is a classifier (or more exactly, a class probability estimator):

Classifier: $p_\theta(z = 1|\boldsymbol{x})$ ,

That estimates the probability of a sequence, $\boldsymbol{x}$, being a member of an optimal set, $z = 1[\boldsymbol{x} \in \mathcal{S}]$, where $1[\cdot]$ is an indicator function that evaluates to 1 if the condition is true, and 0 otherwise. $\theta$ are the parameters of the classifier. Here we define the set as the sequence having a lesser aggregated rank than a threshold, $\mathcal{S} = \{\boldsymbol{x} : R(\boldsymbol{x}) < \tau\}$. Equivalently, we can re-write our classifier as estimating:

$$p_\theta(z = 1|\boldsymbol{x}) = p_\theta(R < \tau|\boldsymbol{x}).$$

The final component is the variational generative model, that estimates the distribution of optimal solutions:

Variational generative model: $q_\phi(\boldsymbol{x}) \approx p(\boldsymbol{x}|z = 1) = p(\boldsymbol{x}|R < \tau)$,

Where again $\phi$ are the parameters of the generative model. This final conditional generative model is learned using a variational inference objective that uses the classifier and prior models to trade-off generating new sequences that have optimal *predicted* properties (*R*) – from the classifier, against being similar to *known* "good" sequences – from the prior. We chose $\tau = 0.5022$ which is the 10$^{th}$ percentile of *R* with SD1 read count > 1, that is the best 10% of our 30801 training sequences are labeled positive. Since we were interested in styrene oxide production, we filtered out variants that did not have more than one styrene dark read. The general VSD training procedure is as follows:

1. Train the classifier, $p_\theta(R < \tau|\boldsymbol{x})$, using 5-fold cross validation on all microfluidics data $z = R < \tau$ and $\boldsymbol{x}$ (using log-loss).
2. Train the prior, $\boldsymbol{x}$, to generate the 30801 sequences, $\boldsymbol{x}$, that had more than 1 read count for SD1 set (using maximum likelihood, irrespective of EF, SD or R).
3. Train the variational generative model, $q_\phi(\boldsymbol{x})$, using the classifier and prior from steps (1) and (2) and the variational inference objective.
4. Sample $s$ sequences from the variational generative model, $\boldsymbol{x}^{(s)} \sim q_\phi(\cdot)$, for testing experimentally.

We visualize this process in Figure 4B. We use the same 2-level convolutional neural network (CNN) architecture for the classifier as in the original VSD paper,[16] and cross validated log-loss results are shown in Figure S2C.

For this work we trialed the two auto-regressive generative backbones used in the original VSD paper[16]; a causal transformer[35], and a long short-term memory (LSTM) recurrent neural network (RNN)[36]. Neither of these generative backbones were constrained to generate mutations only in the 10 targeted sites. Since we had a relatively small budget for experimentally validating the generated sequences, we only chose one of these generative models for the final experiments. To select between the two backbones, we measured the Spearman correlation between the products of 10 direct lead variants, as well as wildtype AbrUPO, and their scores under the generative models. The scores were computed as the average log-likelihood of the variant sequences under the generative models,

$$SCORE_{model} = \frac{1}{N}\sum_{i=1}^{N} \log q_\phi(\boldsymbol{x}_i).$$

We obtained the Spearman correlations in Table S3 – we can see that the LSTM has a much higher correlation with the experimental data than the transformer. We suspect this is because the transformer backbone was overfitting compared to the LSTM – as we can see when looking at a histogram of mutations per site in the generated sequences in. Hence, we chose the LSTM backbone for the G-series variant selection task.

For the final variant generation, we used the VSD-LSTM model to generate 10,000 variants. We then removed any variants that appeared in the original microfluidics dataset, leaving 4,374 novel variants. For the final G-series selection, we ranked these variants by their log-likelihood under the LSTM,

$$G - score(\boldsymbol{x}) \ = \ log\ q_\phi(\boldsymbol{x})$$

then we took the top 5 ranked, and then 5 logarithmically spaced. See Table S5 for the final G-series selection.

### Correlation with pretrained models

Five pretrained models were used to predict the activity of the 21 experimentally validated sequences, comprising the wildtype AbrUPO sequence, 10 R-series variants, and 10 G-series variants. The following pretrained models were used: NetSolP (ESM1b ensemble model), CLIPZyme, ProSmith ESP, SELFprot-Full and the Boltz-2 affinity module.[37–41]

NetSolP predicts solubility and usability (expression*solubility) from a protein sequence using a finetuned protein-language model. CLIPZyme takes a protein sequence and structure and an enzymatic reaction and predicts the probability of the given protein catalyzing the reaction. Protein structures, substrates and products are encoded as graphs and the model is trained with a contrastive learning objective. ProSmith ESP predicts the probability that a small molecule is the substrate for a given protein. Protein and chemical language model encodings are combined into a joint representation and used as input into a transformer network and final gradient boosting model. SELFProt predicts the substrate turnover rate ($K_{cat}$) and Michaelis constant ($K_M$) of a given enzyme-substrate pair. Similar to ProSmith ESP, SELFProt uses a combined protein/small-molecule embedding generated from pretrained protein and chemical language models. The Boltz-2 affinity module uses outputs from the Boltz-2 structure prediction model to calculate the likelihood of a small molecule binding to a given target (probability score) and the affinity of the binding interaction (affinity score). For UPO predictions, Boltz-2 was run using 3 diffusion samples with physics potentials enabled. The heme group and coordinating magnesium ion were included in all predictions and contact constraints were specified to position these cofactors in the correct orientation. For each enzyme/substrate pair, predictions were run twice- once without any constraints on the substrate and once with the substrate constrained to the active site.

The performance of pretrained models was assessed by calculating the Spearman correlation between model scores and experimental results. The permutation method was used to calculate the p-value for each correlation statistic due to the small dataset.

**Table S1.** Clones built for validation of uHTS.

| UPO Variant | Mutations |
|---|---|
| 1 | L84E_G182Y_L231K |
| 2 | L84K_D129N_F179V_L231S |
| 3 | L84Q_G182Y_L231K |
| 4 | L28A_H65T_D129K_L231T |
| 5 | D129N_G182L |
| 6 | L28C_H65C_D129Q_D177S |
| 7 | F8T_L28C_H65K_F179E_A186C_L231C |
| 8 | L84A_A186I |
| 9 | L84R_A186L_L231E |
| 10 | H65R_D129H_D177E |
| 11 | F8A_L28G_H65T |
| 12 | F8V_L28M_H65V_D129M_D177M |
| 13 | F8I_L28M_H65L_D129F |
| 14 | F8M_L28K_D177W_F179R_G182R_A186T |
| 15 | F8P_L28T_H65F_D129F_D177H |
| 16 | F8T_L28P_H65Y_D129Q_D177V |
| 17 | L28I_H65K_F179I_A186V_L231M |
| 18 | F8M_L28K_D177W_F179R_G182R_A186T_L231S |
| 19 | F8C_H65L_D177V |
| 20 | F8K_D129G_F179C_L231K |

**Table S2.** Variant counts for each metric.

| Target | Variant count |
|---|---|
| EFB1 | 2155 |
| EFB2 | 1590 |
| EFB3 | 2455 |
| SD1 | 30801 |
| Union (EFB1, EFB2, EFB3) | 3846 |
| Union (EFB1, EFB2, EFB3, SD1) | 31242 |

**Table S3.** Spearman correlations of μHTS direct lead validation experimental results and both generative model likelihoods. These correlations were used to choose between the LSTM and Transformer models for final G-series creation.

| Experimental Measurement | **LSTM score** $log\ q_{\phi}(\boldsymbol{x})$ | **Transformer score** $log\ q_{\phi}(\boldsymbol{x})$ |
|---|---|---|
| Phenylacetaldehyde (μM) | **0.8091** | 0.4818 |
| Styrene oxide (μM) | **0.6545** | 0.3000 |

**Table S4.** R selected variants.

| Rank | R-score | Mutations |
|---|---|---|
| 1 | 0.045508 | H65R_L84T |
| 2 | 0.048453 | L28C_H65R_L84S_D129Q_D177S |
| 3 | 0.049920 | F8M_L28E_H65S_D129E |
| 4 | 0.053221 | F8S_L28V_H65P_D129S_G182D |
| 5 | 0.054293 | L84M_A186Y_L231V |
| 103 | 0.108083 | F8M_L28V_L84V_F179K_A186Q_L231A |
| 1000 | 0.288362 | L84Y_G182P |
| 10351 | 0.790611 | L84V_D129E |
| 18011 | 0.918425 | L28R_D129K_G182V |
| 23453 | 0.999580 | F179T_G182R_L231P |

**Table S5.** G selected variants.

| Rank | G-score (LSTM log-likelihood) | Mutations |
|---|---|---|
| 1 | -7.186451 | F8G |
| 2 | -7.386762 | H65C |
| 3 | -7.485139 | L84V_D129N |
| 4 | -7.579278 | L84E_G182I |
| 5 | -7.864715 | L84Q_L231V |
| 127 | -9.614363 | L84E_A186V_L231V |
| 167 | -9.877652 | L84M_G182F_L231V |
| 929 | -12.283222 | F8G_G182E_L231V |
| 1849 | -14.582182 | F8I_H65F_D129A_D177M |
| 2630 | -17.302258 | H65K_F179E_L231K |

**SI data:**

SI Data1.xlsx Ultrahigh throughput screening and sequencing data used to train generative protein models.

SI Data2.csv Validation dataset with clones designed from uHTS results.

SI Data3.csv R-series and G-series dataset and predicted performance using pretrained models.

UPO expression cassette.txt Expression cassette used to integrate UPO enzyme variants into *K. phaffii* host strain.
