## Supplemental Figures S1-S5 for "Generative Machine Learning and Microfluidics uHTS: An Efficient Partnership for Enzyme Engineering"

### Figure S1

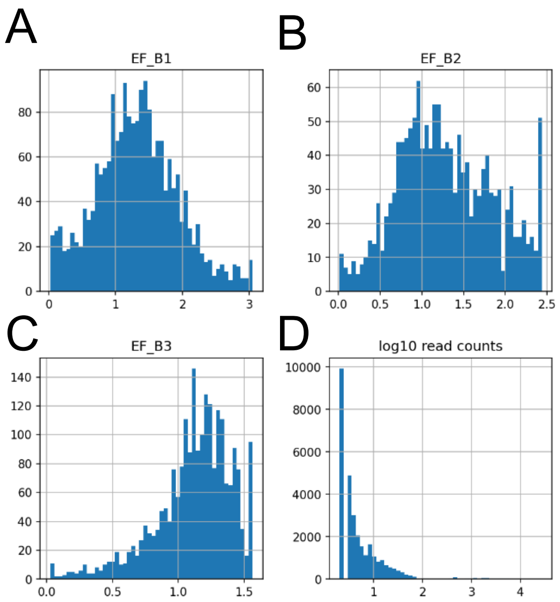

**Histograms of uHTS metrics used for lead selection and generation.** (A-C) Distribution of enrichment factors for variants identified in the three screening thresholds from propylbenzene screening. (D) Distribution of sequencing read counts for variants identified in the dark pool from styrene screening.

### Figure S2

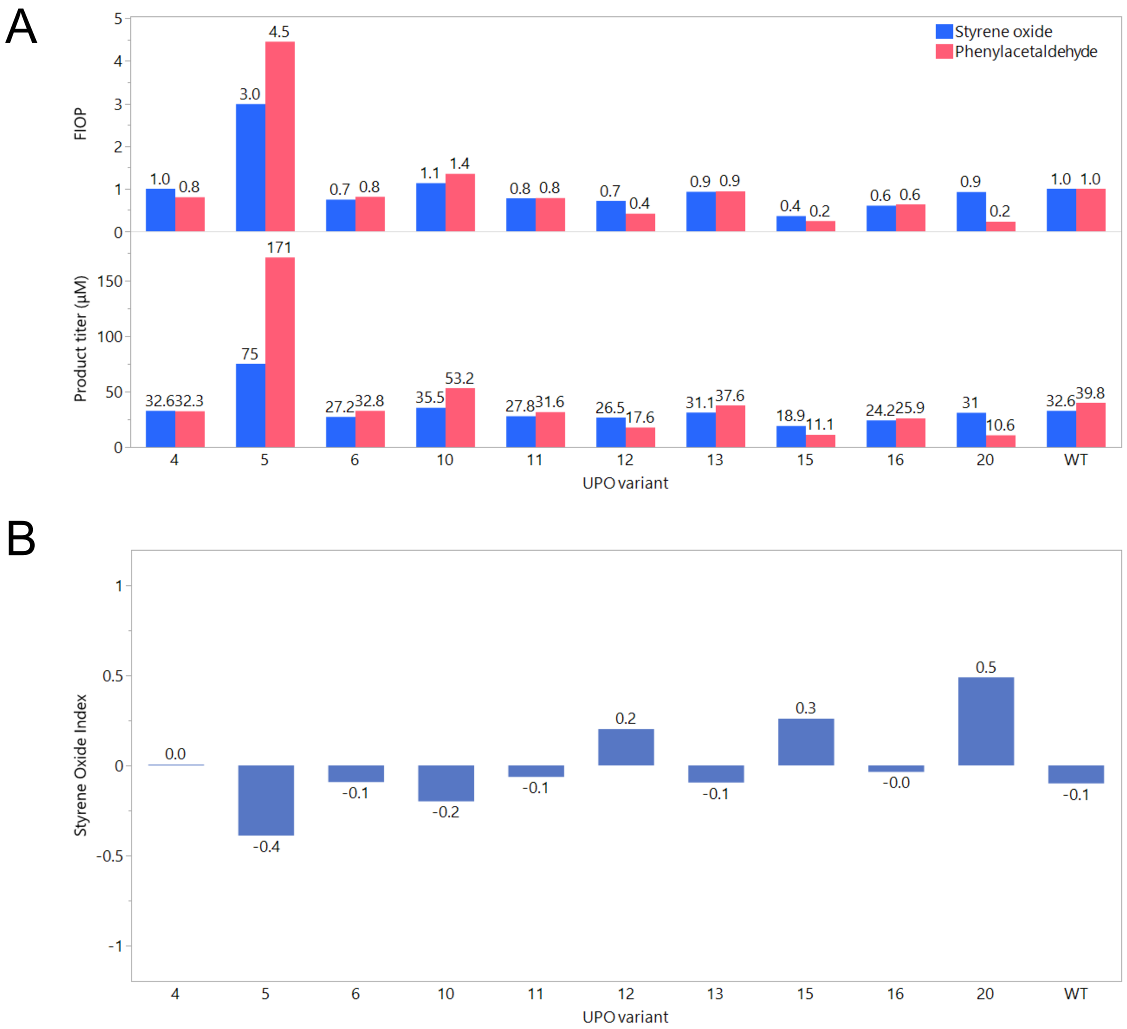

**Ultrahigh throughput direct leads.** (A) Lead AbrUPO variants from uHTS screening were assessed for productivity of styrene oxide and phenylacetaldehyde by gas chromatography. Fold increase over parent (FIOP) was calculated by normalizing product titers of each UPO variant by that of WT UPO. (B) Styrene oxide index (SOI) was calculated for each variant. An SOI of 1 corresponds to formation of only styrene oxide and an index of -1 corresponds to only phenylacetaldehyde.

### Figure S3

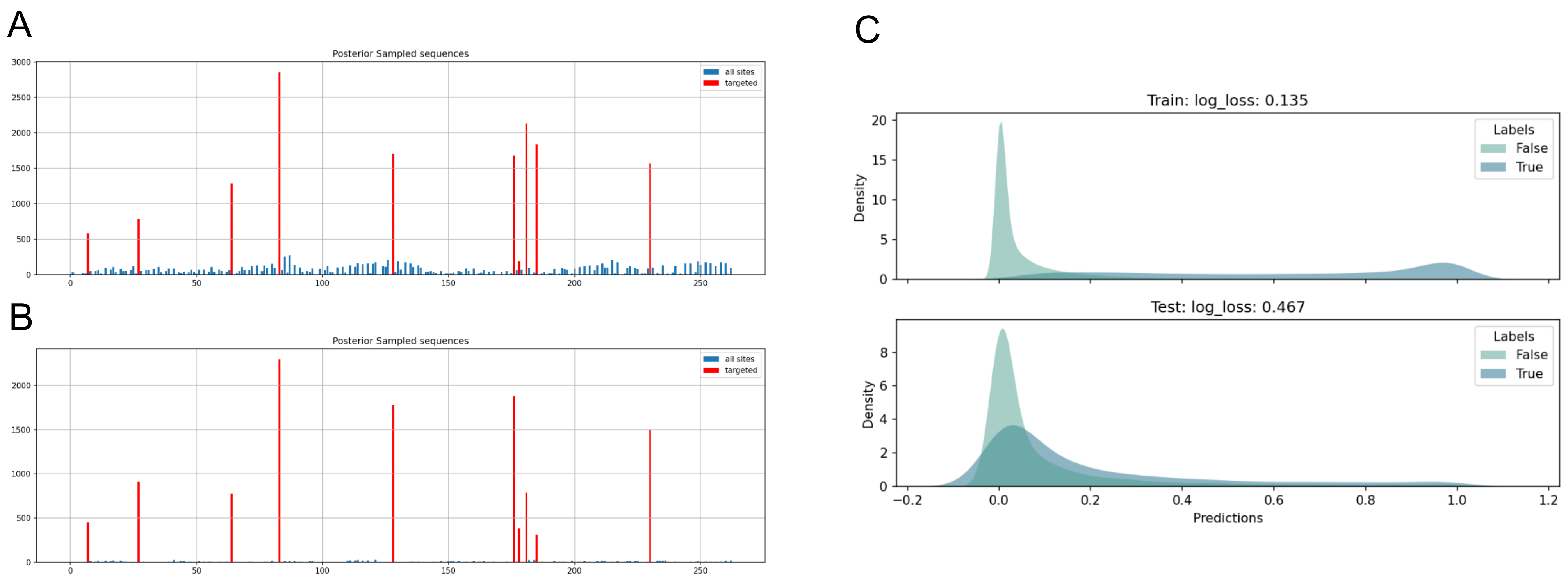

**Model performance evaluations.** Histograms of mutations per site for (A) the VSD LSTM backbone and (B) the VSD transformer backbone. (C) In- and out-of fold cross validation results for the CNN classifier used to train the variational generative model.

### Figure S4A

A

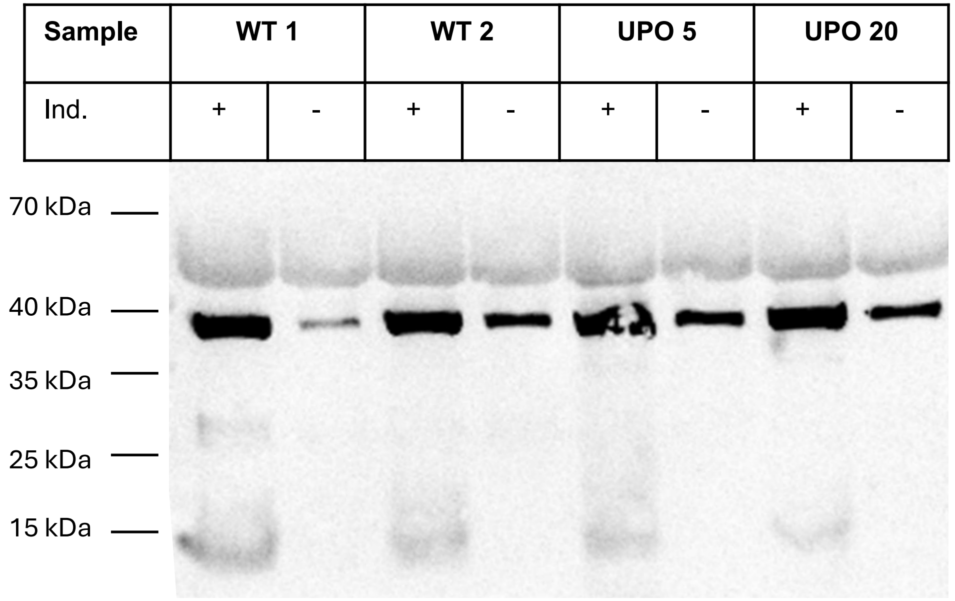

**Western blotting of AbrUPO variants.** AbrUPO variants were built and subjected to western blotting either with or without methanol induction. All samples were subjected to Endo Hf treatment to deglycosylate AbrUPO. Detection utilized an anti-His6 antibody. Expected size of AbrUPO is 30 kDa. Induction-dependent bands were observed at ~38 kDa, ~30 kDa, and ~15 kDa. (A) Two clones of wildtype AbrUPO, UPO 5, and UPO 20.

### Figure S4B

B

| **Sample** | **R1** | | **R2** | | **R3** | | **R4** | | **R5** | | **R103** | | **R1000** | | **R10351** | | **R18011** | | **R23453** |
|---|---|---|---|---|---|---|---|---|---|---|---|---|---|---|---|---|---|---|---|
| Ind. | + | - | + | - | + | - | + | - | | | | | | | | | | | |

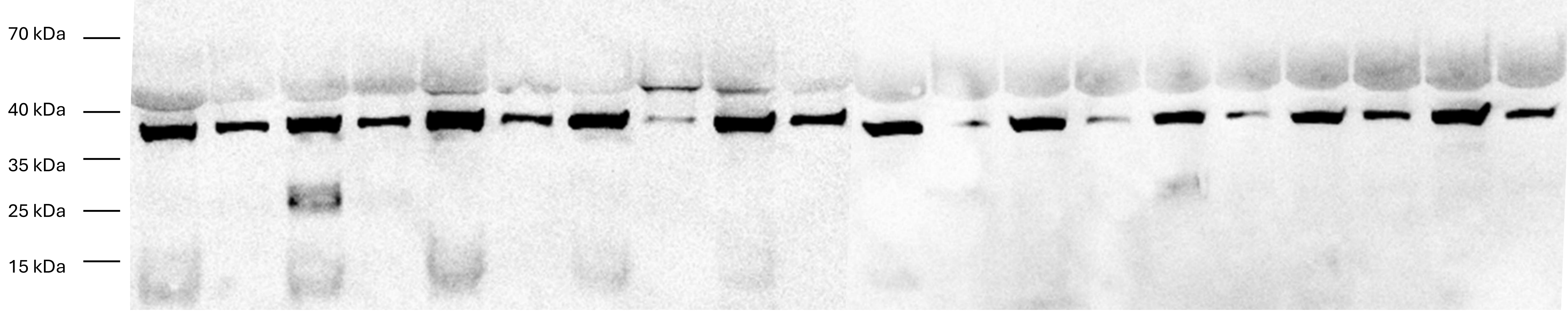

(B) The R-series UPO variants.

### Figure S4C

C

| Sample | G1 | | G2 | | G3 | | G4 | | G5 | | G127 | | G167 | | G929 | | G1849 | | G2630 | |
|---|---|---|---|---|---|---|---|---|---|---|---|---|---|---|---|---|---|---|---|---|
| Ind. | + | - | + | - | + | - | + | - | + | - | + | - | + | - | + | - | + | - | + | - |

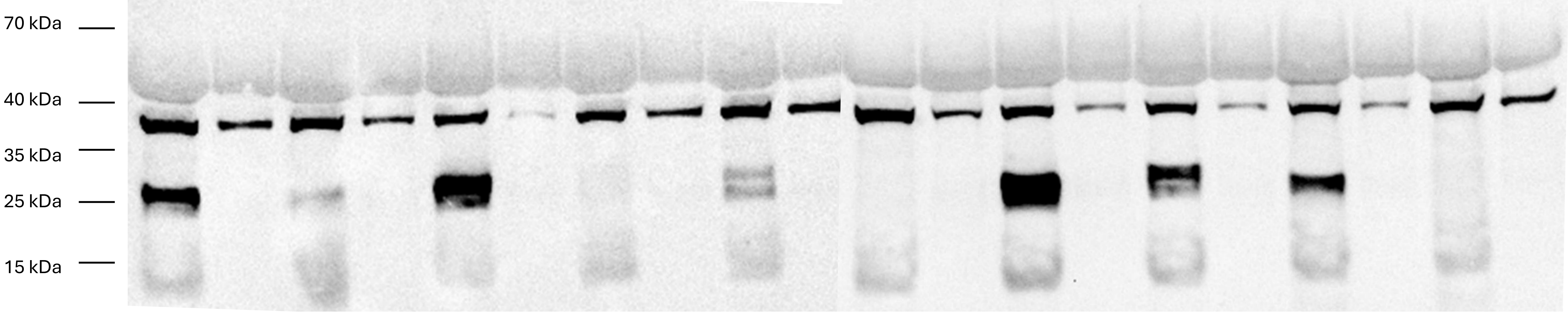

(C) The G-series UPO variants.

### Figure S5

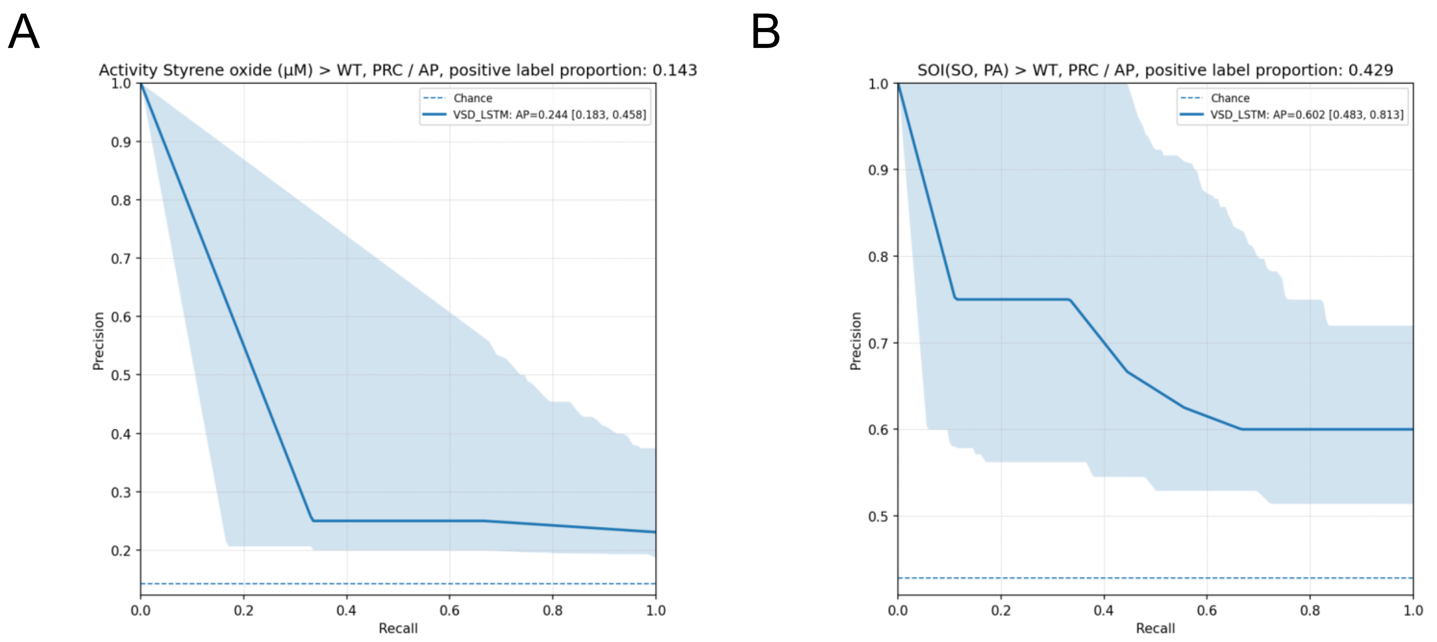

**Bootstrapped precision-recall (PR) curves of the VSD LSTM model's selection performance.** (A) PR curve for higher styrene oxide production than wildtype. (B) PR curve for higher styrene oxide index (SOI) compared to wildtype. The proportion of positive labels is in the figure titles. AP means "average precision", and it is computed as the average precision for all recall values in the curves. AP = 1 would be a perfect predictor, and AP = Chance means the predictor would be no better than random.
